# Treatment with a selective histone deacetylase (HDAC) 1 and 2 inhibitor in aged mice rejuvenates multiple organ systems

**DOI:** 10.1101/2023.08.29.555280

**Authors:** Alessandra Tammaro, Eileen G. Daniels, Iman M. Hu, Kelly C. ‘t Hart, Kim Reid, Rio P. Juni, Loes M. Butter, Goutham Vasam, Rashmi Kamble, Aldo Jongejan, Richard I. Aviv, Joris J.T.H. Roelofs, Eleonora Aronica, Reinier A. Boon, Keir J. Menzies, Riekelt H. Houtkooper, Georges E. Janssens

**Author notes:** Correspondence: R.H. Houtkooper, G.E. Janssens. Equal contribution (Alessandra Tammaro, Eileen G. Daniels, Iman M. Hu).

## Abstract

The process of aging increases the risk of developing age-related diseases, which come at great societal healthcare costs and suffering to individuals. Meanwhile, targeting the basic mechanisms of aging can reduce the risk of developing age-related diseases during aging, essentially resulting in a ‘healthy aging’ process. Multiple aging pathways exist, which over past decades have systematically been confirmed through gene knockout or overexpression studies in mammals and the ability to increase healthy lifespan. In this work, we perform transcriptome-based drug screening to identify small molecules that mimic the transcriptional profiles of long-lived genetic interventions in mammals. We identify one small molecule whose transcriptional effects mimic diverse known genetic longevity interventions: compound 60 (Cmpd60), which is a selective inhibitor of histone deacetylase 1 (HDAC1) and 2 (HDAC2). In line with this, in a battery of molecular, phenotypic, and bioinformatic analyses, in multiple disease cell and animal models, we find that Cmpd60 treatment rejuvenates multiple organ systems. These included the kidney, brain, and heart. In renal aging, Cmpd60 reduced partial epithelial-mesenchymal transition (EMT) *in vitro* and decreased fibrosis *in vivo*. For the aging brain, Cmpd60 reduced dementia-related gene expression *in vivo*, effects that were recapitulated when treating the APPSWE-1349 Alzheimer mouse. In cardiac aging, Cmpd60 treatment activated favorable developmental gene expression *in vivo* and in line with this, improved ventricular cardiomyocyte contraction and relaxation in a cell model of cardiac hypertrophy. Our work establishes that a systemic, two-week treatment with an HDAC1/2 inhibitor serves as a multi-tissue, healthy aging intervention in mammals. This holds potential for translation towards therapeutics that promote healthy aging in humans.

## INTRODUCTION

Increased age in individuals is linked to increased age-related chronic disease^1^. Although aging was long considered a passive process, it is now recognized that the rate of aging can be regulated by so-called longevity pathways^2^. These pathways are diverse, and can include modulation of the insulin-signaling pathway (e.g. targeting insulin-like growth factor 1 (IGF1)^3^, insulin-like growth factor 1 receptor (IGF1R)^4^, or insulin receptor substrate 1 (INSR)^5^), by modulation of mitochondrial biology (e.g. overexpression of sirtuin 6 (SIRT6)^6^), or by improving DNA repair (e.g. overexpression of the mitotic checkpoint gene BUB1, improving genomic stability^7^). Accordingly, many of these genetic interventions influence defined hallmarks of aging, including genomic instability, telomere attrition, epigenetic alterations, a loss of proteostasis, deregulated nutrient sensing, mitochondrial dysfunction, cellular senescence, and stem cell exhaustion^2,8^.

The body of evidence demonstrating genetic interventions that modulate healthy longevity offers the potential for pharmaceutical development targeting these pathways, with the hopes to improve health in the elderly. These pharmaceutical interventions, termed geroprotectors, as they protect the gerontological part of life, are increasingly being uncovered^9–12^. In light of this, a first major testing of one of these compounds is underway in humans with the diabetes drug metformin and the ‘treating aging with metformin’ (TAME) clinical trial^13^, to determine the ability to decrease the incidence of age-related diseases in the elderly.

While there are multiple candidate geroprotectors in line for testing in humans^11^, there is still a great need for second-generation geroprotectors that are more potent and better recapitulate the longevity benefits resulting from genetic interventions. To address this need and circumvent the inherent difficulties of screening for such molecules, which require identifying a proper screening marker, assay development, and chemical screening, our team has been pioneering transcriptome-based drug screening for longevity interventions. For example, our approaches have identified HSP90 inhibitors as proteostasis-inducing longevity interventions^12^, identified longevity compounds with minimized probabilities of side effects in humans^14^, identified the acetylcholine receptor as a target to activate the pro-longevity transcription factor FOXO3^15,16^, and the antiretroviral zidovudine to activate the pro-longevity transcription factor ATF4^17^. In addition to our own work, *in silico* drug screening has been used to identify a novel treatment for metabolic disorder^18^, identify mimetics for the calorie restriction longevity intervention^19^, and in general, de-risk early-phase drug screening^20^.

In the current work, we performed multiple *in silico* drug screens using transcriptional profiles of 2837 small molecules testing their ability to mimic known genetic longevity interventions. We identified one compound that was most commonly found to mimic the transcriptional profile of the genetic longevity interventions. This benzamide-based small molecule, termed compound 60 (Cmpd60; aka Merck60 or BRD 692), is a selective histone deacetylase 1 and 2 (HDAC1/2) inhibitor. Cmpd60 was previously shown to repress growth in certain hematologic malignancies *in vitro*^21^ and *in vivo* to cross the blood-brain barrier to reduce anxiety in mice^22^. Here, we used a combination of molecular, phenotypic, and bioinformatic analyses in multiple disease cell models and mouse models for age-related disease to establish if Cmpd60 acts as a geroprotector. Indeed, we found that Cmpd60 treatment rejuvenates multiple organ systems, including the kidney, brain, and heart. This is in line with our finding that Cmpd60’s transcriptional signature mimics diverse longevity interventions, and with other individual accounts of certain (pan or class-specific) HDAC inhibitors benefiting individual diseases^23^. Our work establishes for the first time specifically HDAC1/2 inhibition as a healthy aging intervention in mammals, further demonstrates that a single molecule can have pleiotropic beneficial effects for healthy aging on multiple organ systems, and paves the way for the development of more potent geroprotective therapeutics in mammals capable of recapitulating the benefits of diverse known genetic longevity interventions.

## RESULTS

### *In silico* transcriptome screening for pharmaceuticals mimicking genetic longevity interventions

In order to identify small molecules that could recapitulate the benefits of multiple genetic longevity interventions, we consulted the GeneAge database, where we found 75 genetic interventions (i.e. either knockdown/outs or overexpressions), which have been documented to extend lifespan^24^. We next turned to the library of integrated network-based cellular signatures (LINCS), an online database and software suite containing mRNA signatures of both drug-treated and genetically perturbed human cell lines^25,26^. Cross-referencing our list of 75 genetic longevity interventions with genetic perturbation cell lines, we found transcriptional signatures were available in the LINCS database for 25 of these (Figure 1A). These 25 interventions, along with FOXO3 overexpression as recently described^15^, were used to query the LINCS database consisting of high-certain transcriptomes of 2837 small molecules present in 8 core cell lines (PC3, VCAP, A375, HA1E, HCC515, HT29, MCF7, and HEPG2), and identify those whose transcriptomic signatures were most similar to at least one of the genetic longevity intervention’s transcriptomes (LINCS score > 90). To ensure the highest likelihood that our drug list would indeed benefit the aging process, we imposed a filter on the query, requiring that a drug’s known target must also be included as a genetic perturbation hit. This resulted in 498 compounds mimicking at least one genetic longevity intervention (Figure 1A). Finally, drugs were ranked according to the number of genetic longevity interventions they transcriptionally mimicked, to form a prioritization ranking (Figure 1B).

When exploring the ranked list of drugs mimicking the most longevity interventions, we noted many well studied drugs in the context of aging (Supplemental Figure 1). For example, the 3^rd^ ranking drug, mimicking 10 out of the 25 genetic interventions, was sirolimus, well-known to extend lifespan in diverse model organisms^27^. Furthermore, ranked 4^th^ and 5^th^ included other molecules that extend lifespan in *C. elegans*, including digoxin^28^, taxifolin^29^, genestein^30^, and catechin^31^. Indeed, many top ranked small molecules from our screen either extend lifespan in model organisms, or have other direct links to age-related pathways (Supplemental Figure 1). However, the top ranked compound, which is the only one to bear transcriptional similarity to 12 out of the 25 genetic longevity interventions, was termed ‘‘compound 60’ (Cmpd60, or ‘Merck60’), and had not yet been explored in the context of healthy aging (Figure 1B).

Cmpd60 is a benzamide-based small molecule that selectively inhibits histone deacetylase 1 and 2 (HDAC1/2). Interestingly, Cmpd60 mimicked the effects of the metabolic related genetic longevity interventions including knockdown of AKT (AKT^KD^), and knockdown of multiple components of the insulin signalling pathway including INSR^KD^ and IRS1^KD^ (Figure 1B), in line with reports that pan-HDAC inhbition can prevent insulin resistance and obestiy in mice fed a high fat diet^32^. Furthermore, since HDAC inhibitors as a drug class may harbor some of the most promising geroprotective compounds^23^, we believed Cmpd60 was an intruiging molecule to further explore in the context of geroprotection.

**Figure 1.**
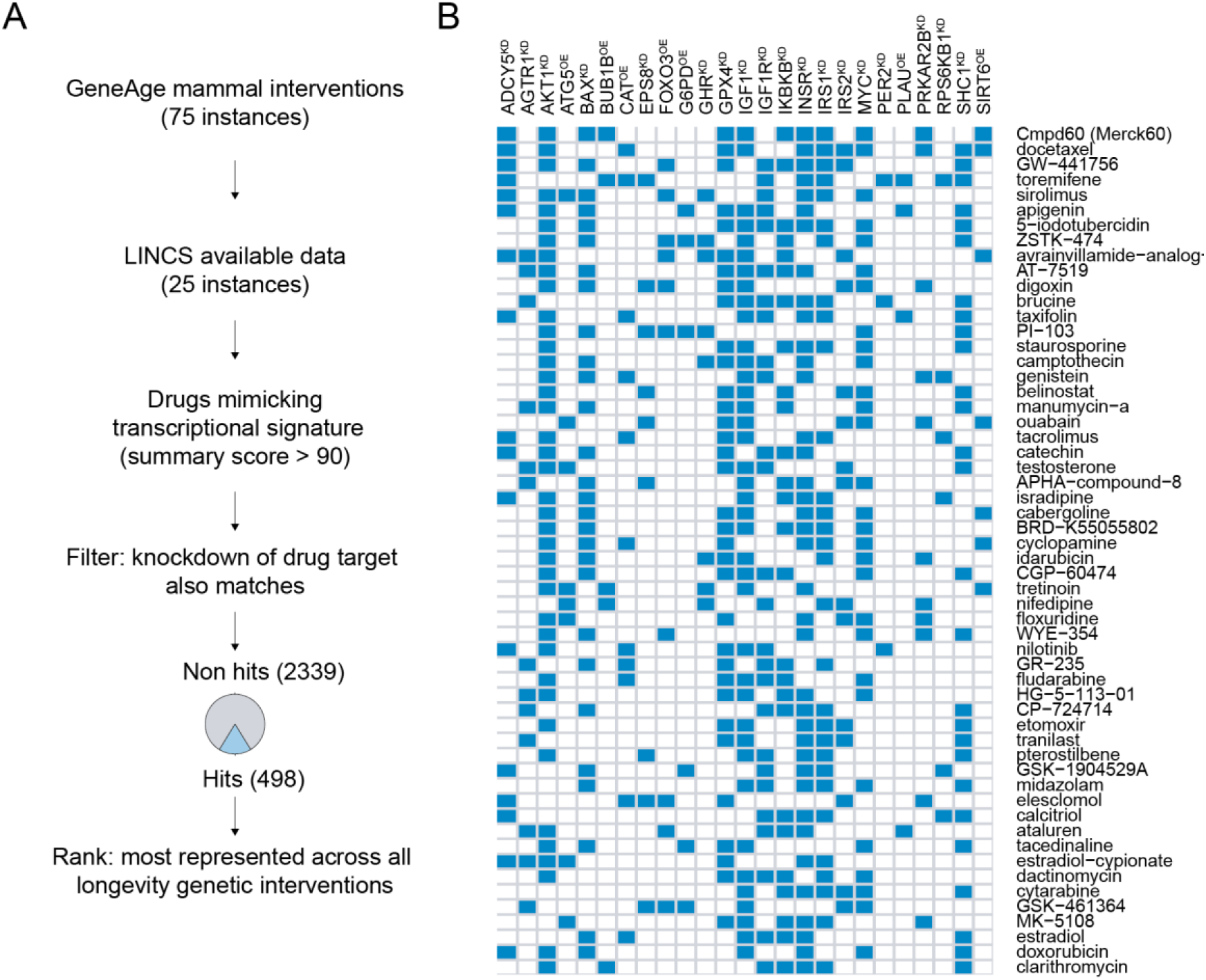
Compound screen strategy and result to identify geroprotectors mimicking genetic longevity interventions. A) General outline of screening strategy. GeneAge database was consulted for listing of genetic interventions, which were cross referenced against the LINCS transcriptome database of cellular purturbations. Compounds best matching a genetic longevity intevention at the transcriptional level were selected for further evaulation, and only those whose drug targets were also present in knockdowns in the screen were included. This resulted in 498 compounds which were ranked based on how many different genetic longevity interventions their transcriptional profiles could recapitulate. B) The top drugs ranked first (top, Cmpd60 as top-ranked small molecule), to last (bottom) according to how many genetic interventions they mimick (blue indicates a positive hit).

### Cmpd60 in aged mice restores youthful molecular and physiological renal parameters

In order to investigate Cmpd60’s potential protective effects during aging, we first turned to an *in vitro* model of renal fibrosis, a hallmark of age-related kidney disease. During aging, senescent tubular epithelial cells (TECs) accumulate in the kidney^33^, which produce a wide range of profibrotic mediators, such as transforming growth factor-beta (TGF-β)^34^. This profibrotic cytokine in turn affects TECs’ phenotype, promoting a partial epithelial-mesenchymal transition (EMT), ultimately leading to renal fibrosis^35^. Partial EMT in TECs is marked by an increased expression of the mesenchymal gene Alpha Smooth Muscle actin (αSMA) and a decrease in Zonula occludens-1 (ZO-1) and E-cadherin^36,37^. Indeed, treating TECs with recombinant TGFβ was sufficient to significantly increase αSMA and reduce ZO-1 protein expression (Figure 2A). We then tested if Cmpd60 could prevent EMT in TECs. We used a dose of 1μM Cmpd60, which is a non-toxic dose (Supplemental Figure 2A) that effectively increases histone acetylation levels at histone H3K18 and H4K8 (Supplemental Figure 2B-C). Strikingly, Cmpd60 partially prevented EMT upon TGFβ stimulation, reducing αSMA and increasing ZO-1 and E-cadherin protein expression (Figure 2A and supplemental Figure 2D).

To determine Cmpd60’s geroprotective effects on the kidney at the molecular and physiological levels, we proceeded to treat aged male mice (20 months old) for 14 days with either Cmpd60 (22.5mg/kg) or control (Figure 2B). This dosing regimen was based in part on previous studies with Cmpd60^22^. EchoMRI measurements showed that fat mass, lean mass, and total body weight did not change between treated and untreated mice suggesting that Cmpd60 was tolerated at the dose used (Supplemental Figure 2E), which matched the observation that blood biochemistry markers for renal and liver toxicity did not differ between the two groups (Supplemental Figure 2F). Assessing acetylation levels revealed an increase of H4K8 acetylation in Cmpd60 treated kidneys, demonstrating efficacy of the intervention (Figure 2C, Supplemental Figure 2G).

To assess the molecular effects of Cmpd60 on the kidney, we performed RNAseq transcriptomics on kidneys from the treated and untreated aged mice (Supplemental Table 1). Samples could be readily differentiated using partial least squares discriminant analysis (PLS-DA) (Figure 2D). Exploring the data further, we calculated differential expression between the groups, where we noted that inhibition of HDAC1/2 with Cmpd60 imparted clear differences on the transcriptional landscape (p<0.01, Supplemental Table 1). To better understand what these changes were, we performed gene ontology (GO) term and KEGG pathway analyses on the up and down regulated genes (Figure 2E, Supplemental Table 2). Here we found the top upregulated go term was one often associated to longevity and healthy aging, namely, glutathione metabolic processes^38^ (Figure 2E). These genes included Glutathione S-Transferase genes (*Gstm1, Gsta3*, and *Gsta4*) (Supplemental Figure 2H), an important family of detoxifying and cytoprotective enzymes crucial for longevity^39^ and as protective mechanism against the development of renal fibrosis^38,40–42^. Given the decrease in partial EMT observed in the *in vitro* model, we sought to assess how these molecular changes manifest themselves at the physiological level. We performed histological analysis of renal fibrosis by analyzing collagen content detected with picro sirius red. Markedly, we found that the aged Cmpd60 treated mice showed less age-related renal fibrosis than their untreated counterparts (Figure 2F-G). Taken together, these findings suggest Cmpd60 alters the transcriptional landscape in aged kidney cells, shifting it towards a profile protective from oxidative stress and conducive to a reduction of renal EMT and age-related kidney fibrosis.

**Figure 2.**
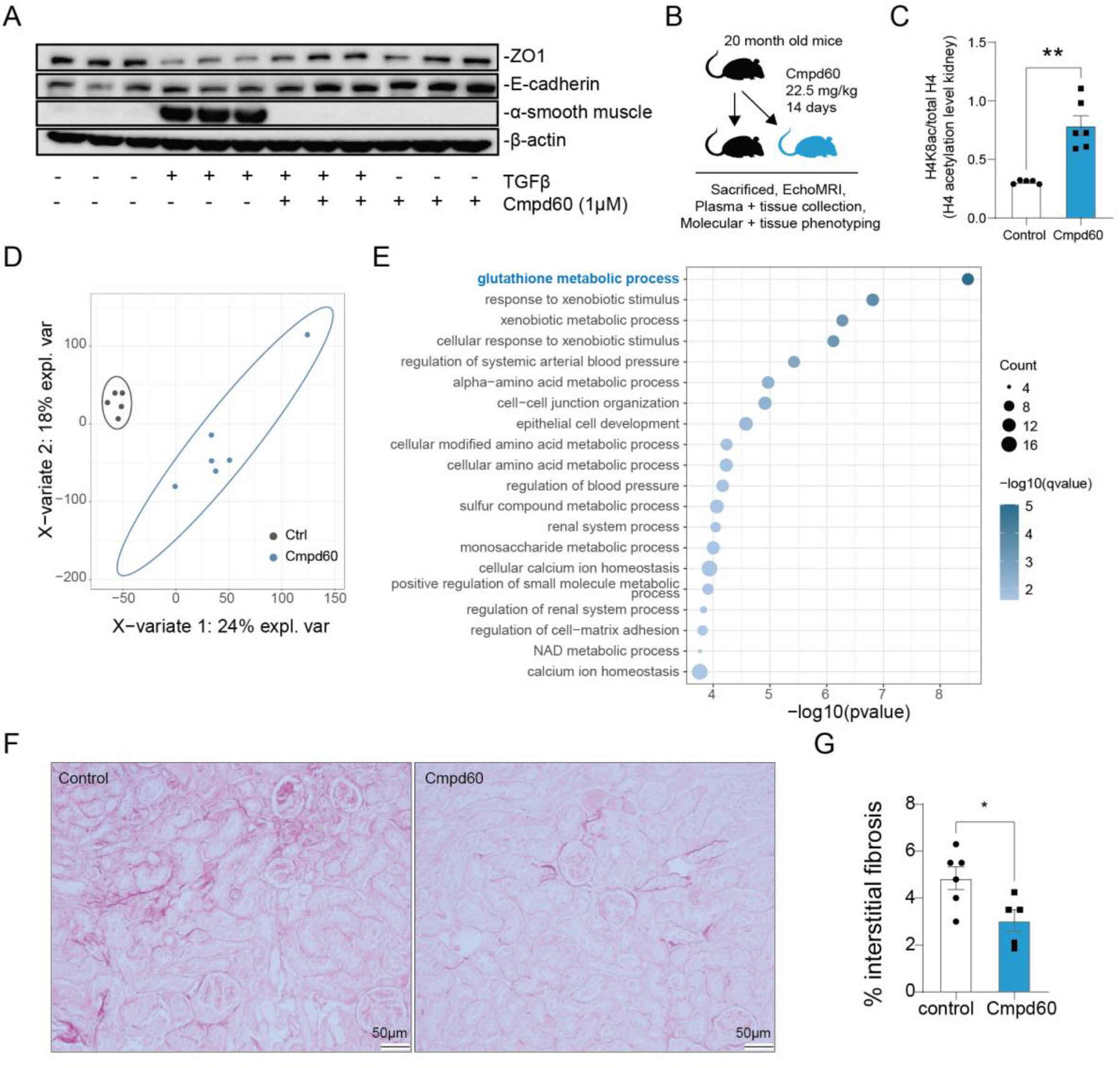
Influence of Cmpd60 on aging kidney. A) Representative western blot of Tubular epithelial cells (TECs) treated with 20ng/ml of recombinant TGFβ and with and without Cmpd60 (1 μM) for 72hrs. Protein lysates of TECs blotted for anti-ZO-1, anti-E-cadherin, anti-αSMA and β-actin. Cmpd60 suppresses markers for partial EMT, a hallmark of age-related renal fibrosis (n=3)/group. B) Schematic of aged mouse treatment regimen with Cmpd60 and analyses. C) Relative Histone H4 acetylation levels (H4K8Ac) assessed by western blot in renal tissue of aged mice with and without Cmpd60 treatment. Protein expression was normalized against H4 total and expressed as mean ±SEM. Mann Whitney test was used to determine statistical differences, **P<0.01. (n=5-6)/group. D) PLS-DA analysis of aged mice treated with and without Cmpd60 (n=5-6)/group. E) Top GO terms of upregulated processes in aged mice treated with Cmpd60 (see also Supplemental Table 2). F) Representative histological images of Picro Sirius Red staining in kidney of aged mice with and without Cmp60. (n=5-6)/group. G) Quantification of interstitial fibrosis determined by the percentage of positive Picro Sirius Red staining/high power field, in mice treated with and without Cmpd60. Percentage of positive staining was assessed with Image J software. Data are expressed as mean ±SEM and the Mann Whitney test was used to determine statistical significance, *P<0.05, (n=5-6)/group.

### Cmpd60 treatment protects against detrimental brain aging processes

Having seen clear benefits of Cmpd60 treatment to the aged renal system, and noting prior work of others that demonstrated Cmpd60’s ability to cross the blood brain barrier^22^, we inquired the effects of Cmpd60 on the aged brain. Assessing histone modification in the brain revealed increased acetylation levels (Figure 3A, Supplemental Figure 3A). Establishing this, we proceeded to perform RNAseq transcriptomics on brains of treated and untreated aged mice (Supplemental Table 3). Here, PLS-DA readily separated the two groups (Figure 3B), and we applied the same cutoff as for the kidney to assess differential expression (P<0.01, Supplemental Figure 3B). Interestingly, assessing enriched GO terms and KEGG pathways revealed an alteration in oxidative phosphorylation processes, down regulated upon treatment (Figure 3C, Supplemental Table 4). Remarkably, the KEGG pathway of Alzheimer’s was also downregulated upon Cmpd60 treatment (Figure 3C). This included genes also involved in oxidative phosphorylation such as the NADH:Ubiquinone Oxidoreductase Subunits (NDUFs) (Figure 3D), in line with the finding that decreasing mitochondrial capacity can reduce amyloid-β toxicity^43^.

Observing this potential beneficial effect, we next asked if Cmpd60 treatment could help prevent neurological decline in a dementia model. To address this, we turned to the APPSWE-1349 mouse model, a transgenic mouse overexpressing an isoform of human Alzheimer beta-amyloid (βA), which shows clear signs of impaired spatial referencing at 9-10 months of age^44^. We proceeded to treat APPSWE-1349 mice and control littermates for 14 days with either Cmpd60 (22.5mg/kg) or control (Figure 3E). We used mice younger than those that show full physiological symptoms, aged 6-7 months, to ensure the greatest chance of intervening in the early, molecular-based processes that occur and contribute to βA accumulation and neurodegeneration. Likewise we focused on molecular readouts to assess efficacy. Performing RNAseq transcriptomics on brain of these mice (Supplemental Table 5) and PLS-DA, revealed a strong separation of the non-treated transgenic mice, but less separation of the Cmpd60 treated transgenic mice from the control littermate mice (Figure 3F). This suggested Cmpd60 treatment was shifting the transgenic mouse profile away from a disease profile towards a non-disease profile. Comparing the differential gene expression between the treated and untreated transgenic mice (Supplemental Figure 3C) and performing GO term and KEGG pathway enrichments (Supplemental Table 6), revealed that Cmpd60 reduced ribosomal gene expression (Supplemental Figure 3D), while increasing membrane potential, ion transport, and cognitive processes (Supplemental Figure 3E), changes previously reported to be conducive to decreased dementia risk^45–47^. Taking into account all four groups, namely the transgenic and control mice, both untreated and treated, allowed for an analysis of gene expression changes that Cmpd60 induced, unique to the transgenic disease model. Here, relevant for Cmpd60’s potential effects in dementia specifically, we found an up regulation of memory related GO terms (Figure 3G, Supplemental Table 6). Some of the differentially expressed genes in this category included *Pla22g6*^48^, *Cx3cr1*^49^, *Ncam1*^50,51^, *Cyfip1*^52^ (Figure 3H), genes whose expression have been shown to benefit cognitive processes.

Finally, to determine how these transcriptional changes may manifest at the physiological level, we performed histological analysis of brains from the transgenic mice, either Cmpd60 treated or untreated. While the mice we studied were younger than the age at which aggregates are clearly visible, we found suggestive evidence that pre-aggregates were less present in Cmpd60 treated mice. Specifically, 4 out of 7 untreated mice showed aggregates (57%), while only 2 out of 6 Cmpd60 treated mice showed aggregates (33%) (Supplemental Figure 3F). Taken together, our findings suggest Cmpd60 modifies the brain transcriptional landscape in a manner protective against the brain aging changes and counter to dementia related processes.

**Figure 3.**
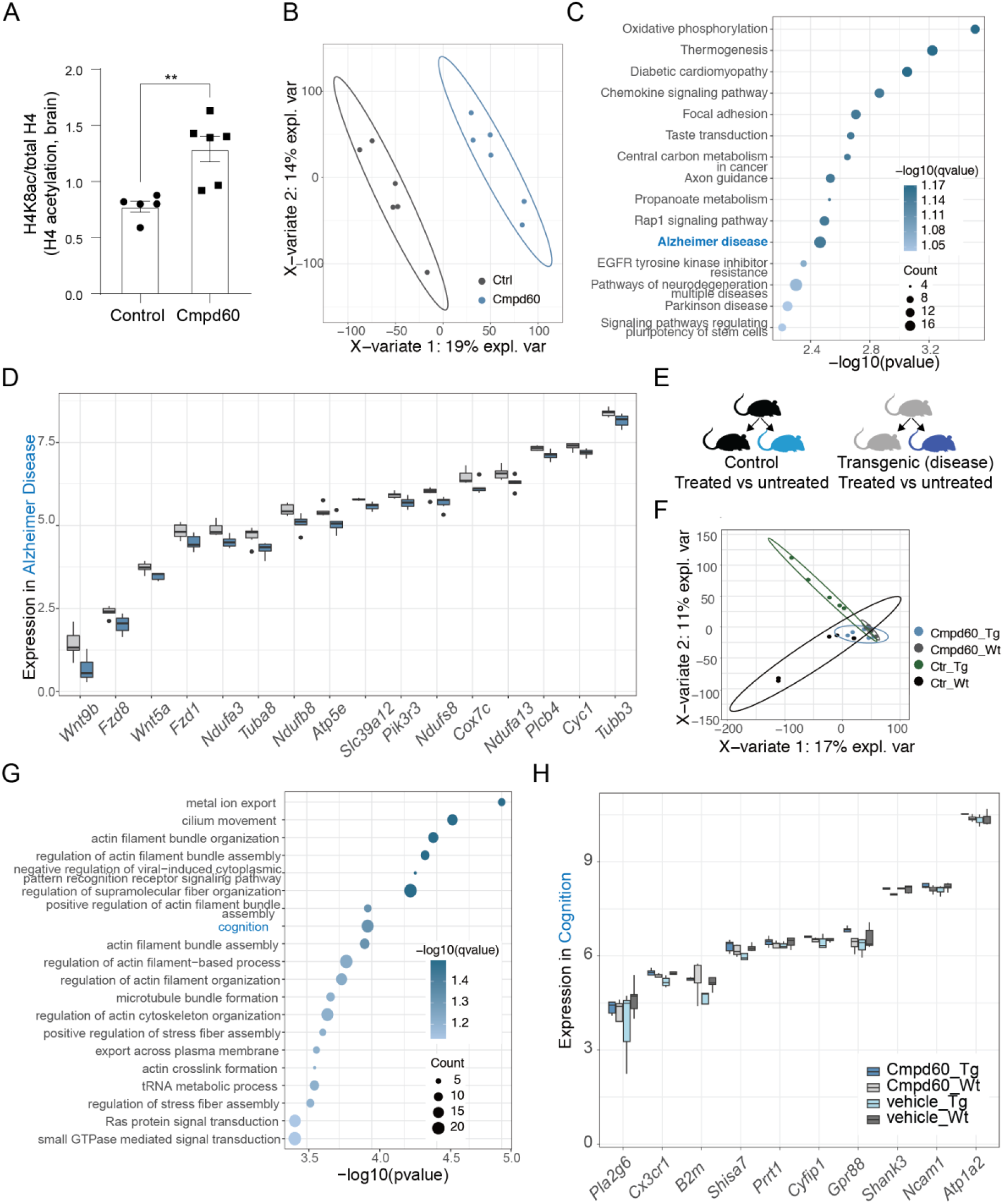
Cmpd60 treatment supports healthy brain aging. A) Relative expression of Histone H4 acetylation levels (H4k8Ac) assessed by western blot in brain tissue of aged mice treated with control and Cmpd60. Protein expression was normalized against H4 total and expressed as mean ±SEM. Mann Whitney t-test was used to determine statistical differences, **P<0.01. n=5-6/group. B) PLS-DA of RNA-seq transcriptome comparing Cmpd60 treated and untreated brain, n = 6 per group. C) Downregulated KEGG terms resulting from Cmpd60 treatment. D) Boxplot of counts per million (CPM) expression values of genes in Cmpd60 treated mouse brain from the GO term enrichment of Alzheimer Disease. Fill represents condition, grey for control and blue for Cmpd60. E) Schematic of dementia mouse model and treatment. F) PLS-DA of RNA-seq transcriptome comparing dementia mice, controls, treated and untreated. G) Up Go terms of interaction between the 4 groups, revealing altered cognitive processes. H) Boxplot of CPM expression values of genes from the GO term enrichment of cognition in (E). Fill represents condition, grey for control and blue for Cmpd60.

### Cmpd60 treatment improves cardiac function

Having noted Cmpd60’s beneficial effects on the aged kidney and brain, with relevance for two serious and under-treated age-related dysfunctions of renal failure and dementia, we next inquired as to the effects of Cmpd60 on one of the organs most contributing to age-related death: the heart. Our initial analysis did not reveal significant acetylation changes in histone H3 or H4 (Supplemental Figure 4A-C). Nonetheless, to further explore Cmpd60’s cardiac-related effects more deeply, we performed RNAseq transcriptomics on hearts from aged treated and untreated mice (Supplemental Table 7). Here we again observed samples to be readily distinguishable upon PLS-DA (Figure 4A).

Upon evaluating differential expression in the Cmpd60 treated versus untreated heart samples, we noted far greater transcriptional changes following Cmpd60 treatment in the heart compared to either the kidney or brain. Accordingly, we applied a stricter cut-off to assess differential expression (adjusted p-value < 0.05) (Figure 4B). Although we found fewer GO enrichments and KEGG pathways related to altered oxidative phosphorylation processes, strikingly, we found the top enriched GO terms were related to heart valve development, suggesting profound changes influencing heart function may be occurring upon Cmpd60 treatment (Figure 4C, Supplemental Table 8). This upregulation included genes such as SMAD Family Member 6 (*Smad6*), ADAM Metallopeptidase With Thrombospondin Type 1 Motif 9 (*Adamts9*), and Elastin Microfibril Interfacer 1 (*Emilin1*), members of gene families who have all been linked to cardiovascular outcomes, with either deficiency proving detrimental or abundance proving beneficial^53–55^ (Figure 4D).

Spurred by these promising findings, we turned to an *in vitro* assay of cardiac functioning. Here, we assessed contraction (percentage of sarcomere shortening) and relaxation (return velocity) in adult rat ventricular cardiomyocytes. Remarkably, and in line with our *in vivo* findings at the transcriptional level, we found Cmpd60 treated ventricular cardiomyocytes showed both an improved contraction and relaxation parameters (Figure 4E-F). Taken together, this suggests Cmpd60 treatment modifies the cardiac transcriptional level and manifests itself at the functional level to improve age-related cardiac outcomes.

**Figure 4.**
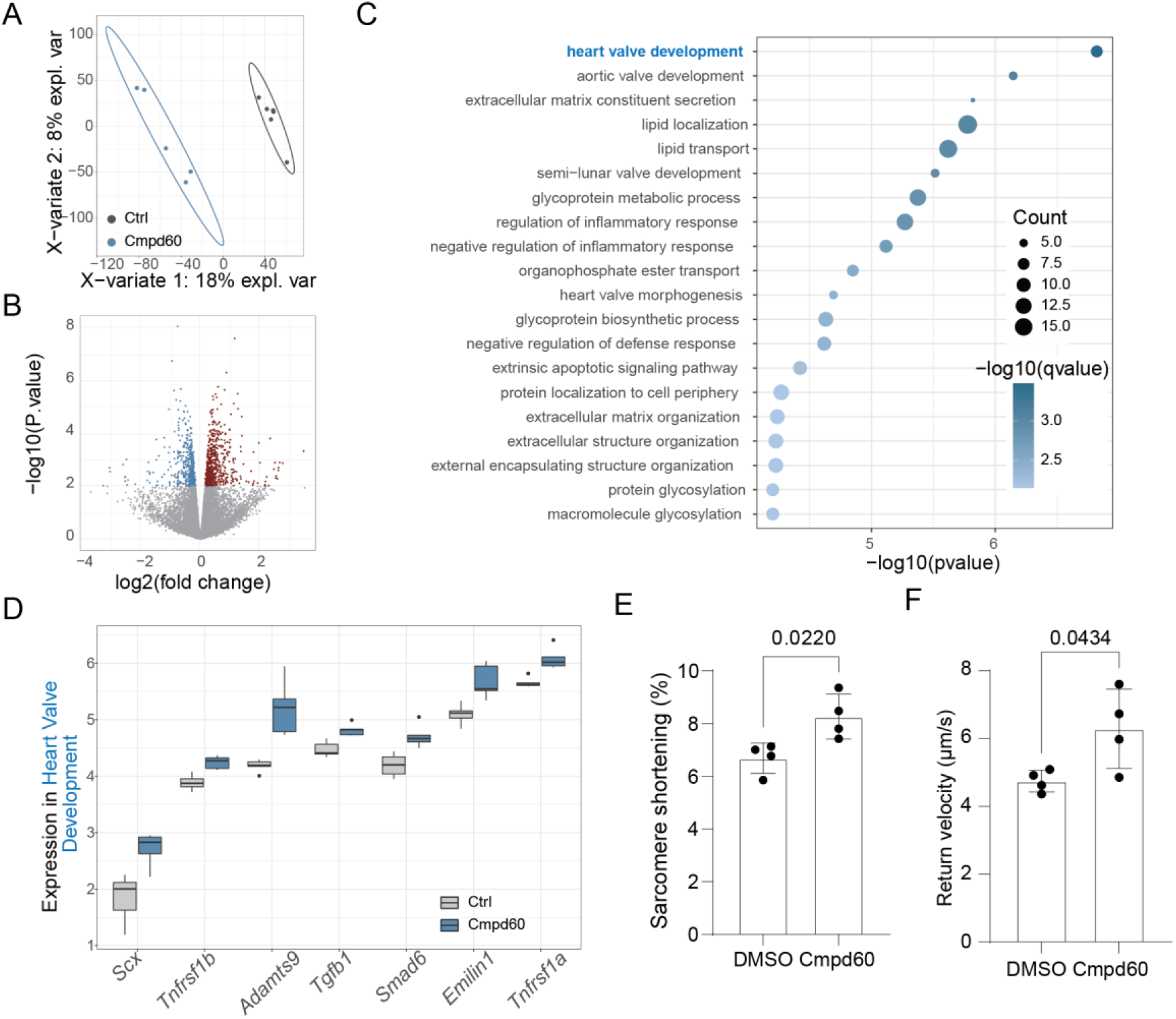
Cmpd60 treatment benefits cardiac tissues. A) PLS-DA of RNAseq, aged heart, treated vs untreated (n = 5-6/group). B) Volcano plot of RNAseq differential expression, aged heart, treated vs untreated (n = 5-6/group). Genes with p-value < 0.01 were colored (red: up-regulated, blue: down-regulated). C) Top GO Terms from upregulated genes. D) Boxplot of CPM expression values of genes in Cmpd60 treated mouse heart from the GO term enrichment of Heart Valve Development. E) Treatment of cardiomyocytes with Cmpd60 increased contraction, as shown by increased % sarcomere shortening. F) Treatment of cardiomyocytes with Cmpd60 improved relaxation, as assessed by higher return velocity (n=4, corresponding to 4 independent experiments; 30-40 CMs were measured per condition per experiment; data are represented as mean ± SD, *p*<0.05, unpaired *t*-test).

### A consensus model of Cmpd60’s effects

Having identified tissue-specific benefits of Cmpd60, we next inquired whether a conserved expression profile existed amongst the different tissues of the treated mice. To accomplish this, we assessed the overlap of differentially expressed genes in the kidney (p-value<0.05), brain (p-value<0.05), or heart (adjusted p-value<0.05). We identified 41 genes upregulated (Figure 5A) and 30 genes downregulated (Figure 5B) in common between the three tissues following Cmpd60 treatment. Amongst these 71 genes, for example, were genes including upregulated *Mapk3, Tgm2*, and *Spns2*, and downregulated *Mrps28* and *Fzd8* (Figure 5C). Transcription factor analysis querying diverse motif databases revealed six motifs (transfac-pro-M00797, cisbp-M6275, swissregulon-hs-HIF1A.p2, transfac-pro-M00466, transfac-pro-M07043, homer-TACGTGCV-HIF-1a) associated with Hif1a target genes (Figure 5D), suggesting Cmpd60 treatment increases oxidative stress resistance, an observation in line with the main transcriptional changes observed in the kidney and brain. Taken together, our work suggests both tissue specific effects of Cmp60 treatment, such as *Gsta2/3/4* and *Gstp1/3* in the kidney, *Wnt5a* in the brain, and *Scx* and *Emilin1* in the heart, as well as common transcriptional changes shared between tissues, oriented around *Hif1a* target gene expression. Together, the cumulated effects of these molecular changes may result in the age-reversing qualities we observed following Cmpd60 treatment in old mice.

**Figure 5.**
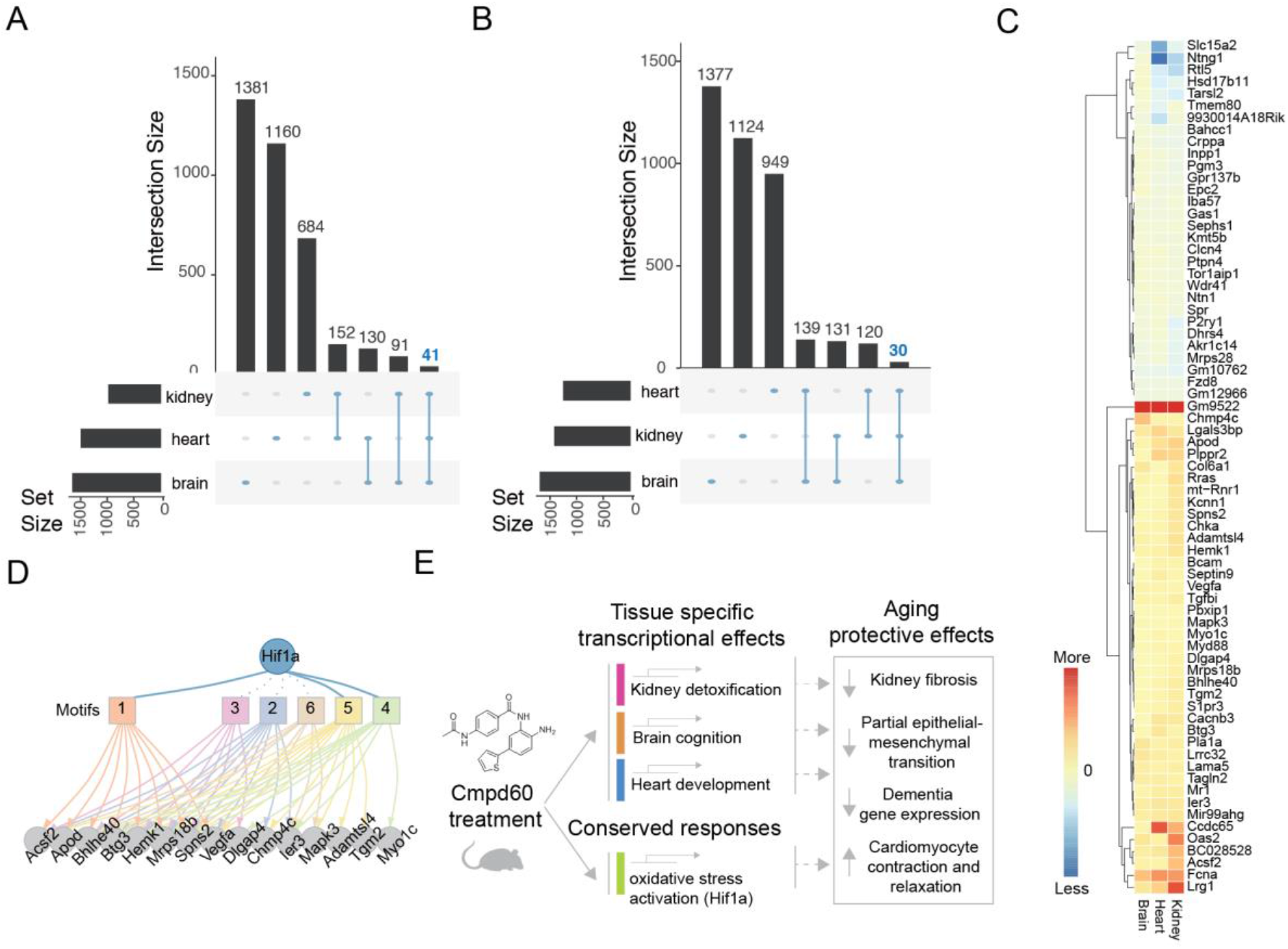
A consensus model of Cmpd60’s effects. A) Comparison of the unique and shared upregulated genes in the three tissues; kidney, heart and brain. 41 genes are commonly upregulated in the three tissues (highlighted in blue) B) Comparison of the unique and shared downregulated genes in the three tissues. 30 genes are commonly downregulated in the three tissues (highlighted in blue) C) Heatmap of the log fold change of genes with shared regulation in three tissues (for visualization purposes, log fold changes exceeding 2 were capped at 2, while values below -2 were capped at -2). D) Network for transcription factor Hif1a, one of the top predicted TFs based on motif overrepresentation of the commonly changed genes among the three tissues. Squares represents different motifs annotated to Hif1a. Edges connect each motifs to the genes contributing to its enrichment. E) Model of Cmpd60’s geroprotective effects, which are due to both tissue specific and conserved transcriptional changes, producing net aging-protective effects.

## DISCUSSION

In this work, we used an *in silico* drug screening platform and identified a single molecule, the HDAC1 and HDAC2 inhibitor Cmpd60, which possessed transcriptional signatures mimicking diverse genetic longevity interventions. In line with this, Cmpd60 demonstrated distinct rejuvenating effects across multiple organ systems. In the kidney, Cmpd60 treatment increased protective gene expression related to oxidative stress regulation and reduced fibrosis possibly via reduced partial EMT detected in *in vitro* studies. In the brain, Cmpd60 treatment showed transcriptional changes conducive to improved cognitive functioning and molecular indications of neuroprotection in both naturally aged brain and a dementia mouse model brain. In the heart, Cmpd60 resulted in cardiac remodeling related transcriptional changes and benefitted cardiomyocyte functioning.

With Cmpd60 demonstrating such diverse age-related benefits across multiple organs, a question remains as to how these effects are mediated. HDAC inhibition has previously been suggested to benefit health through a plethora of mechanisms, including FOXO3 activation^56^, Klotho upregulation^57^, or reversing age-related acetylation changes, amongst others^23^. Notably, these have all been explored in diverse models and organs. The likeliest answer therefore is that HDAC inhibition modifies a tissue-specific epigenetic landscape, creating beneficial tissue specific responses (Figure 5E). Since aging is accompanied by alterations in histone acetylation patterns and global loss of transcriptional control^58^, one tantalizing possibility is that Cmpd60 reverses these acetylation changes to reinstate a more youthful transcriptional profile in a tissue specific manner. Our findings at the transcriptional level, including an upregulation of oxidative stress protection and alterations in metabolic gene expression, catered to each organ, support this idea. These manifested themselves as a rejuvenation of multiple organ systems at the cellular and physiological levels. It remains to be seen how each organ achieved such benefits, and how these findings can further translate to benefit human health.

As most studies on HDAC inhibitors focus on one specific tissue, our study is unique in that it looks at the effects of HDAC inhibition in three different organs; kidney, brain and heart. This enabled us to recognize an overlapping gene expression profile in all three tissues; associated with *Hif1a* target genes. Although we identified tissue-specific benefits of Cmpd60, HDAC inhibitors are also known for their undesirable side effects^59^. Despite of, or thanks to, their many diverse on- and off-target effects, HDAC inhibitors nonetheless benefit a range of preclinical age-related disease models^23^. We therefore recommend future research to assess dose-dependent effects of HDAC1/HDAC2 inhibitors in multiple organs.

## Acknowledgements

AT is financially supported by the NWO-FAPESP joint grant on healthy ageing, executed by ZonMw (no. 457002002). GEJ is supported by a VENI grant from ZonMw, and AGEM Talent and Development grants. RHH is financially supported by an ERC Starting grant (no. 638290), a VIDI grant from ZonMw (no. 91715305) and by the Velux Stiftung (no. 1063). EGD is supported by a ZonMw travel grant (no. 446001023). MH is supported by a China Scholarship Council 2020 grant. EA is supported by, EU H2020-EpiEpiNet (No 952455).

## Competing interests

The authors declare no competing interests.

## Contributions

A.T., E.G.D., R.H.H. and G.E.J. conceived and designed the project. E.G.D. K.R., K.J.M. and R.I.A. organized and performed animal husbandry and Cmpd60 treatment and physiological animal measures. A.T. and L.M.B. performed kidney in vitro studies and with R.K. performed all biochemical analyses. A.T., E.A. and J.J.T.H.R. performed pathology analyses. A.J. and G.V. performed bioinformatics analyses. I.M.H. performed mouse transcriptomic analyses. K.C.H. R.P.J. and R.A.B. designed cardiac experiments and interpretations. A.T., E.G.D., R.H.H. and G.E.J. wrote the manuscript with contributions from all authors.

## Methods

### *In silico* compound screen

The online library of integrated network-based cellular signatures (LINCS)^25,26^ was accessed in September 2017 through the cloud-based software platform CLUE (https://clue.io/) as previously described^60^. The ‘touchstone’ core dataset consisting of transcriptome signatures of eight cell lines (PC3, VCAP, A375, HA1E, HCC515, HT29, MCF7, HEPG2) of 2837 different small molecule treatments, 3799 different gene knock-downs, and 2160 different gene overexpressions was used. From the gene knock-downs or overexpression datasets, those genetic longevity interventions known to confer lifespan extension in mouse models were used, (accessed in 2017 from GeneAge, filtering for significant, positive lifespan effects in mice^24^). Individual queries were performed for each genetic longevity intervention, producing lists of compounds with similar transcriptional signatures. Compound lists were ranked and included a summary score consolidating cell line data, ranging from −100 (opposing the genetic longevity signature) to 100 (mimicking the genetic longevity signature). These were downloaded as.gct files (version 1.3). A cutoff was applied to the ranked list such that compounds with a score > 90 were considered to match the transcriptional signature of a longevity intervention. Drug lists were further filtered, such that a drug was only included as a hit, if its drug target (i.e. the knockdown of the drug target) also passed a summary score cutoff >90 for the genetic longevity intervention in question. The final ranking was producing by tallying the total number of genetic longevity interventions a compound could mimic (theoretical total 25), where more than one compound could reach the same rank. Only one compound reached the top rank (12 out of 25), Cmpd60.

### Cmpd60

Cmpd60, also known as Merck60 with Broad Institute ID number BRD6929 and CAS No.: 849234-64-6 was aquired from ChemShuttle (USA, China), Catalog No.: 151025.

### *In vitro* experiments

Murine Immortalized proximal tubular epithelial cells (TECs) were generated as previously described^61^ and cultured in DMEM/HAM F12 (Gibco) supplemented with 10% fetal calf serum, penicillin/streptomycin, 2mM L-glutamin (Invitrogen), 5μg/ml insulin (Gibco), 5μg/ml transferrin (Gibco), 5ng/ml selenite (Gibco), 40pg/ml Tri-iodo-thyrionine (Sigma), 36ng/ml hydrocortisone (Sigma) and 20ng/ml EGF (Sigma). TECs were maintained in culture at 33°C in medium supplemented with 10ng/ml IFNγ (Prospec) to maintain SV40 expression (ref). One week before experiments were performed, TECs were differentiated at 37°C for 7 days in presence of complete medium without IFNγ. TECs were stimulated with 20ng/ml murine recombinant TGFβ (Prospec) for 72 hours in DMEM/F12 supplemented with 10% fetal calf serum, penicillin/streptomycin and 2mM L-glutamin. Cmpd60 was added either together with TGFβ for 72 hours or added in the last 6 hours of the experiment. After 72hrs cells were washed with PBS and processed for RNA or protein isolation.

### Cell viability assay

TECs were plated in a 96 well plate (Greiner). After 24 hours the cells were stimulated with Cmpd60 at different concentration (0,5-100μM) and incubated for 72 hours. The next day 1mg/ml MTT (Sigma-Aldrich) was added to the medium and the cells were incubated for another 45 minutes at 37°C until formazan crystals became visible. Medium was discarded and DMSO was added to dissolve the crystals. The optical density was measured at 570nm at microplate reader (Clariostar). Cell viability was established in relation to the control cells.

### Cell lysates and immunoblot

RIPA lysis buffer (50mM Tris pH7.5, 0,15M NaCl 2mM EDTA, 1% deoxycholic acid, 1% nonidet P40, 0,1% SDS supplemented with 4mM Na_3_VO_4_, 0,5mM NaF and protease inhibitors (Sigma)) was added to the cells at the end of the experiment. Cells lysates were centrifuged and protein concentration was measured using a BCA assay kit (Thermo Scientific). Twenty μg of protein was loaded onto a 4-12% Bis-Tris gradient gel (Invitrogen) and separated proteins were transferred on PVDF membrane (Millipore). After blocking aspecific signal, membranes were incubated overnight at 4°C with primary antibody. Primary antibodies used for in vitro experiments: acetylated Histone H3 (Lys 18 Cell signaling #9675), total Histone 3 (Cell signaling #9715), acetylated Histone H4 (Lys8, Cell signaling #2594), total Histone 4 (Cell Signaling #2592), a-SMA (Dako M085101) ZO-1 (Invitrogen #617300), E-cadherin (Cell signaling #3195) and β-actin (Millipore). The following day membranes were incubated with horseradish peroxidase (HRP)-conjugated secondary antibodies (anti rabbit-HRP (Dako), anti msIgG2a-HRP (Southern Biotech) & anti msIgG1-HRP (Southern Biotech)) for 1 hour at RT. Detection was done by ECL western blotting substrate (Thermo Scientific) and images were obtained on a LAS 4000 (ImageQuant). Band intensity was quantified through ImageJ.

### Tissue lysates and immunoblot

Freeze dried tissues (kidney, brain and heart) were homogenized in lysisbuffer (120mM Tris pH 6.8, 4% SDS, 20% glycerol supplemented with protease inhibitors) and stored overnight at -20C. The next day the homogenates were passed through a 21G needle and protein was measured using a BCA kit (Thermo Scientific). Twenty μg of protein was loaded on a 4-12% Bis-Tris gradient gel (Invitrogen) and separated proteins were transferred on PVDF membrane (Millipore). After blocking aspecific signal, membranes were incubated overnight at 4°C with primary antibody. The following primary antibodies were used for immunoblot: Histone 3 (Lys 18 Cell signaling #9675), total Histone 3 (Cell signaling #9715), acetylated Histone H4 (Lys8, Cell signaling #2594), total Histone 4 (Abcam ab7311), b-actin (Abcam) and GAPDH (Cell Signaling #2118). The following day membranes were incubated with horseradish peroxidase (HRP)-conjugated secondary antibodies (anti rabbit-HRP (Dako) and anti msIgG1-HRP (Southern Biotech)) for 1 hour at RT. Detection was done by ECL western blotting substrate (Thermo Scientific) and images were obtained on a LAS 4000 (ImageQuant). Band intensity was quantified through ImageJ.

### Plasma biochemical analysis

Renal, liver and body toxicity were determined by measuring plasma levels of urea, creatinine, Aspartate aminotransferase (ASAT), Alanine transaminase (ALAT) and Lactate Dehydrogenase (LDH). These parameters were determined by enzyme reactions using standard autoanalyzer methods by our hospital research services.

### Histology and Immunostaining

Paraffin-embedded kidney and brain tissues were processed for (immuno)histological analysis. To quantify the percentage of interstitial fibrosis, Picro Sirius red histological staining was performed to detect collagen content. Kidney tissue slides were incubated with 0.2% Picro Sirius Red (PSR) solution (pH 2.0) for 1h followed by incubation with 0.01M HCl. The amount of PSR-positive staining per high power field (20x magnification) was quantified by Image J software. Beta amyloid plaques in brain slides were identified with beta Amyloid (1-42) antibody (Genetex: GTX134510). Quantification of the percentage of amyloid plaques was performed by the neuropathologist in a blinded manner.

### Animal husbandry and Cmpd60 treatment

#### Aged mice

Natural aged, approx. 20 months old, male BL6 mice were acquired from Taconic and single housed under a 12:12-hour light-dark cycle in a room set to 23°C (+/-0.2°C). All animals were fed a regular chow diet. Mice received daily i.p. injections with either Cmpd60 (n=6) or vehicle (n=7) for 14 days total. Body weight was monitored every 5 days to adjust i.p. volumes to body weight. Cmpd60 treated animals received a dose of 22.5 mg/kg with an i.p. volume of 7.5 ml/kg. Cmpd60 was dissolved in 2% DMSO, 49% PEG400, and 49% saline solution (= vehicle) resulting in a 3 mg/ml concentration.

#### APPSWE-1349 mice

The APPSWE-1349 mice^62^ (BL6 background) were acquired from Taconic. The transgenic mice possess a transgene coding for the 695-amino acid isoform of human Alzheimer β-amyloid (Aβ) precursor protein carrying the Swedish mutation. Animals (∼6-7 months old) were single housed under a 12:12-hour light-dark cycle in a room set to 23°C (+/-0.2°C). All animals were fed a regular chow diet. Mice received daily i.p. injections with either Cmpd60 (n=12) or vehicle (n=12) for 14 days total. Body weight was monitored every 5 days to adjust i.p. volumes to body weight. Cmpd60 treated animals received a dose of 22.5 mg/kg with an i.p. volume of 7.5 ml/kg. Cmpd60 was dissolved in 2% DMSO, 49% PEG400, and 49% saline solution (= vehicle) resulting in a 3 mg/ml concentration.

### Mouse measures and tissue collection

On day 15, following 14 days of Cmpd60 treatment, mice (total body mass) were weighed and loaded into the Echo-MRI (EchoMRI-700 Analyzer) using an A100 antenna insert and then whole-body fat and lean mass were measured. Mice then underwent an overnight fast (10 hours) and then were euthanized using CO_2_, total body weight was determined, followed by exsanguination through cardiac puncture. Blood (600-1000ml) was collected by cardiac puncture into a heparin-coated syringes. Blood samples were centrifuged for 20 min at 4°C and the separated plasma was stored at -80°C for further analysis. Tissues and organs were collected, weighed and either snap frozen in liquid nitrogen or submerged in O.C.T. (Fisher Scientific) and stored at -80°C.

### RNA sequencing: Isolation of mRNA

Mouse tissues were homogenized with a 5 mm steel bead using a TissueLyser II (QIAGEN) for 5 min at frequency of 30 times/second. RNA was extracted according to the instructions of the RNaesy Mini Kit (QIAGEN). Contaminating genomic DNA was removed using RNase-Free DNase (QIAGEN). RNA was quantified with a NanoDrop 2000 spectrophotometer (Thermo Scientific; Breda, The Netherlands) and stored at -80°C until use.

### RNA sequencing: Library Preparation

RNA libraries were prepared and sequenced with the Illumina platform by Genome Scan (Leiden, The Netherlands). The NEBNext Ultra II Directional RNA Library Prep Kit for Illumina was used to process the sample(s). The sample preparation was performed according to the protocol “NEBNext Ultra II Directional RNA Library Prep Kit for Illumina” (NEB #E7760S/L). Briefly, mRNA was isolated from total RNA using the oligo-dT magnetic beads. After fragmentation of the mRNA, cDNA synthesis was performed. This was used for ligation with the sequencing adapters and PCR amplification of the resulting product. The quality and yield after sample preparation was measured with the Fragment Analyzer. The size of the resulting products was consistent with the expected size distribution (a broad peak between 300-500 bp). Clustering and DNA sequencing using the NovaSeq6000 was performed according to manufacturer’s protocols. A concentration of 1.1 nM of DNA was used. NovaSeq control software NCS v1.6 was used.

### RNA sequencing: Read Mapping and Statistical Analyses

Reads were subjected to quality control FastQC^63^ trimmed using Trimmomatic v0.32 (Bolger et al., 2014) and aligned using HISAT2 v2.1.0 (Kim et al., 2015). Counts were obtained using HTSeq (v0.11.0, default parameters) (Anders et al., 2015) using the corresponding GTF taking into account the directions of the reads. Statistical analyses were performed using the edgeR v3.26.8 (Robinson et al., 2010) and limma/voom v 3.40.6 ^68^ R packages. All genes with more than 2 counts in at least 3 of the samples were kept. Count data were transformed to log2-counts per million (logCPM), normalized by applying the trimmed mean of M-values method ^67^ and precision weighted using voom (Law et. al., 2014). Differential expression was assessed using an empirical Bayes moderated t test within limma’s linear model framework including the precision weights estimated by voom ^68,69^. Resulting *p* values were corrected for multiple testing using the Benjamini-Hochberg false discovery rate. Data processing was performed using R v3.6.1 and Bioconductor v3.9. Partial least-squares discriminant analysis (PLS-DA) was performed using mixomics (Rohart et al., 2017) setting a variable of importance (VIP) score of greater than 1 as significant. Resulting *p* values (where applicable) were corrected for multiple testing using the Benjamini-Hochberg false discovery rate. Genes were re-annotated using biomaRt using the Ensembl genome databases (v91).

### Transcriptome analysis and visualization

Data processing was performed using R version 4.1.1. Genes were reannotated using the Ensembl genome database and the biomaRt package^71^. Resulting p-values were corrected for multiple testing using the Benjamini–Hochberg false discovery rate where applicable. Biological process (BP) overrepresentation analysis was performed using Clusterprofiler 4.0.5^72^ and org.Mm.eg.db (version 3.13.0). Gene selection (for figures 2E, 3C, 3G, 4C, supplementary figure 3D, 3E) was done based on p-value less than 0.01, and log Fold change larger than 0 for up-regulated genes, smaller than 0 for down-regulated genes. Partial least squares discriminant analysis (PLS-DA) was performed on normalized cpm value (genes with zero expression were filtered out) using MixOmics version 6.16.3^70^. Upset plots were generated using UpsetR version 1.4.0^73^. For transcription factor (TF) binding motif over-represention, analysis was performed using RcisTarget 1.14.0^74^. Shared up-regulated genes (pvalue < 0.05, log Fold change larger than 0) between brain, kidney and heart were used as input gene list. The same was performed for shared down-regulated genes. The following file (mm9-500bp-upstream-7species.mc9nr.feather) was used to specify the gene-motif rankings. “motifAnnotations_mgi_v9”was used for motif annotation to transcription factors. Additionally, the pheatmap (1.0.12), igraph (1.30) and ggplot2 (version 3.4.2) packages were used to generate heatmaps and various visualizations using colors from RcolorBrewer^75–77^.

### Adult rat ventricular cardiomyocyte isolation and contractility measurement

The animal experiments were performed in accordance with the guidelines from the Directive 2010/63/EU of the European Parliament on the protection of animals used for scientific purposes and approved by the ethics committees of Amsterdam University Medical Centers, VUMC location, Amsterdam, the Netherlands. Adult rat left ventricular cardiomyocytes (CMs) were isolated as described previously^78,79^. Briefly, adult wild-type Wistar rats were terminated under anesthesia, followed by chest opening and heart extraction. The heart was cannulated through the aorta and perfused on a Langendorf perfusion set-up with liberase enzyme solution until the tissue was sufficiently digested. The atria and right ventricle were removed and the left ventricle was minced into small pieces and triturated. Subsequently, the cell suspension was filtered and re-suspended in CaCl_2_ buffers of increasing Ca^2+^ concentrations to reach a final concentration of 1mM. The isolated adult CMs were finally re-suspended in plating medium containing Medium 199 (Lonza, BE12-117F), 1% penicillin/streptomycin (Lonza, DE17-602DE) and 5% fetal bovine serum (PAA, A15-101), and seeded on 1% laminin (L2020-1MG, Sigma)-coated plates (24-well format Costar culture plate, Corning, 3524). One hour after plating, the medium was refreshed with maintenance medium containing Medium 199, 1% penicillin/streptomycin and Insulin-Transferrin-Sodium Selenite Supplement (Sigma-Aldrich; insulin, 10 mg l−1; transferrin, 5.5 mg l−1; and selenium 5μgl−1). Subsequently, the cells were stimulated with 5μM Cmpd60 (or corresponding vehicle, DMSO) for 2 hours at 37°C in humidified air with 5% CO_2_. After the stimulation, the contraction and relaxation of the CMs were measured with the MultiCell microscope system (CytoCypher, Amsterdam, the Netherlands) coupled to the Ionoptix high-speed sarcomere length measuring software (Ionoptix LLC, Westwood, Massachusetts). Unloaded intact rat CMs were monitored following field stimulation, and sarcomere shortening was measured and analyzed with the automated, batch analysis software transient analysis tools (Cytosolver, CytoCypher) to determine the contraction and relaxation profiles of the cells.

### Statistical analyses

Statistical analyses were performed using either R (version 4.1.1) or PRISM (9.5.1) and specific tests and corrections for multiple hypothesis testing are listed in either each experiment’s figure legend or corresponding methods section.

## Notes

### Competing Interest Statement

The authors have declared no competing interest.

