## Supplemental figures for "Treatment with a selective histone deacetylase (HDAC) 1 and 2 inhibitor in aged mice rejuvenates multiple organ systems"

| Ranked drug | | Lifespan or aging related effect |
|---|---|---|
| 3 | sirolimus | Extends lifespan in worms/flies/mice (Johnson 2013) |
| 3 | apigenin | Suppresses inflammation (Perrot et al 2017) |
| 4 | digoxin | Extends lifespan in worms (Wang et al 2017) |
| 4 | taxifolin | Extends lifespan in worms (Benedetti 2008) |
| 5 | staurosporine | Extends lifespan in flies (Spindler et al 2012) |
| 5 | camptothecin | Topo I inhibitor induces hormesis (Zhang et al 2015) |
| 5 | genistein | Extends lifespan in worms (Lee et al 2015) |
| 5 | catechin | Extends lifespan in worms (Saul 2009) |
| 6 | tretinoin | Treatment for photo-aged skin (Kligman 1984) |
| 6 | nifedipine | Extends lifespan in rotifers (McTavish et al 1990) |
| 6 | nilotinib | In silico identified geroprotector (Ziehm 2017) |
| 6 | fludarabine | Improves healthspan in rotifers (Snell 2016) |
| 6 | tranilast | Treatment for inflammaging related diseases (Huang 2018) |
| 6 | pterostilbene | Chemically related to resveratrol (Li 2017) |
| 6 | calcitriol | Vitamin D, extends lifespan in worms (Golegaonkar et al 2015) |
| 6 | dactinomycin | In silico identified geroprotector (Barardo et al 2017) |
| 6 | estradiol | Extends lifespan in mice (Harrison et al 2014) |
| 6 | clarithromycin | Extends lifespan in rotifers (Snell 2016) |

**Supplemental Figure 1 | Known effects of top ranked compounds.** Selection of the top ranked list of drugs mimicking the most longevity interventions in the context of aging.

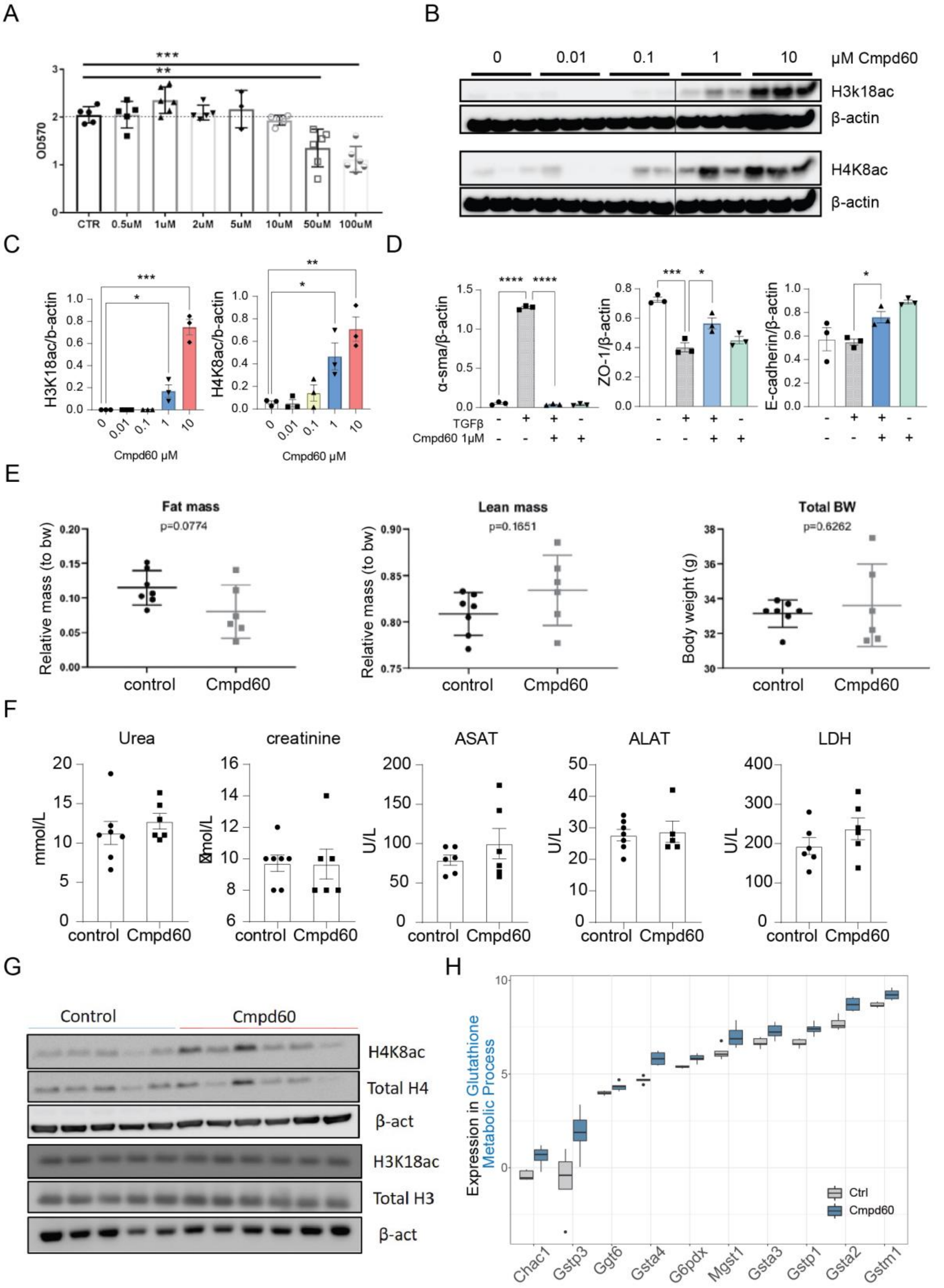


**Supplemental Figure 2 | Cmpd60 and aged kidney**

A) MTT toxicity assay of Cmpd60 at various doses in tubular epithelial cells (TECs) stimulated for 72hours. B) Representative western blot showing expression of H3k18Ac, H4K8Ac and β-actin in TECs stimulated with different doses of Cmpd60 for 72hrs. C) Relative expression of H3k18Ac and

H4K8Ac. Protein expression levels were normalized against β-actin and expressed as mean ±SEM. Three independent experiments were performed. T-test was used to determine statistical differences, *P<0.05; **P<0.01 ***P<0.001. n=3/group. D) Relative expression of α-SMA, ZO-1 and E-cadherin in TECs stimulated with TGFβ (20ng/ml) with and without 1μM Cmpd60. Protein expression levels were normalized against β-actin and expressed as mean ±SEM. Three independent experiments were performed. Student's t-test was used to determine statistical differences, *P<0.05; ***P<0.001; ****P<0.0001 n=3/group. E) EchoMRI of aged mice treated with or without Cmpd60, shows no evidence of toxic effects (Student's t-test). F) Plasma biochemistry markers of aged mice treated with or without Cmpd60, shows no evidence of toxic effects in kidney (urea and creatinine), liver (ASAT and ALAT) and body (LDH) (Student's t-test). G) Representative western blot showing expression of H3K18ac, total H3, H4K8ac, H4 total and β-actin in aged kidneys treated with control or Cmpd60 as indicated in Figure 2B. (n = 5-6/group).
H) Boxplot of counts per million (CPM) expression values of genes in Cmpd60 treated aged mouse kidney from the GO term enrichment of glutathione metabolic process. Color represents condition, grey for control and blue for Cmpd60.

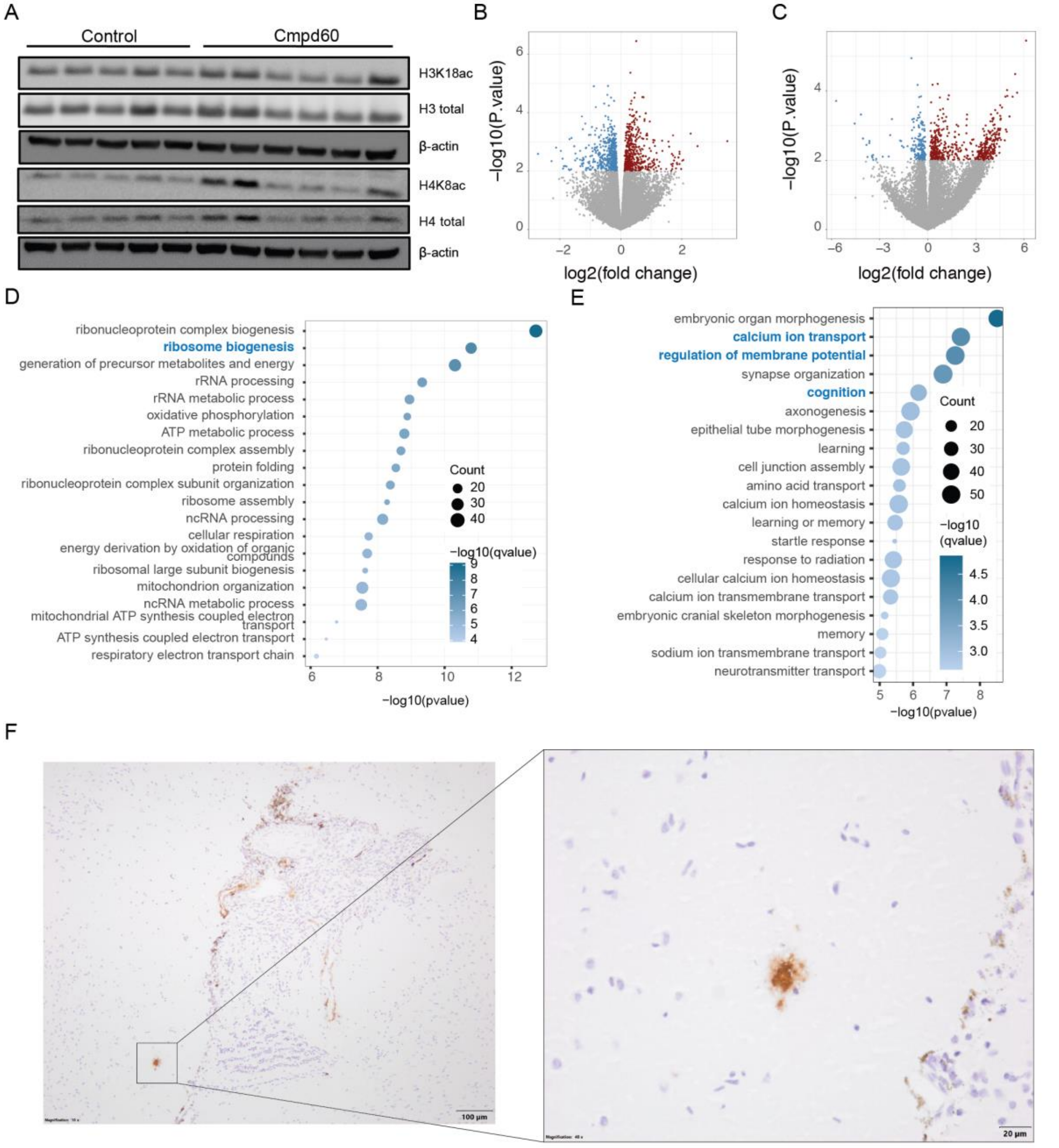


**Supplemental Figure 3 | Cmpd60's effects on the aged and diseased brain**

A) Representative Western blot showing expression of H3K18ac, total H3, H4K8ac, H4 total and β-Actin in aged brains treated with control or Cmpd60 as indicated in Figure 2B. (n=5-6/group). B) Volcano plot of RNAseq differential expression, aged brain, treated vs untreated. Genes with p-value < 0.01 were colored (red: up-regulated, blue: down-regulated). C) Volcano plot of RNAseq differential expression, transgenic brain, treated vs untreated. Genes with p-value < 0.01 were colored (red: up-regulated, blue: down-regulated). D) GO term enrichments DOWN, transgenic brain, treated vs untreated. E) GO term enrichments UP, transgenic brain, treated vs untreated. F) Representative IHC pictures at different magnifications of β-amyloid plaques in transgenic AD mice brain.

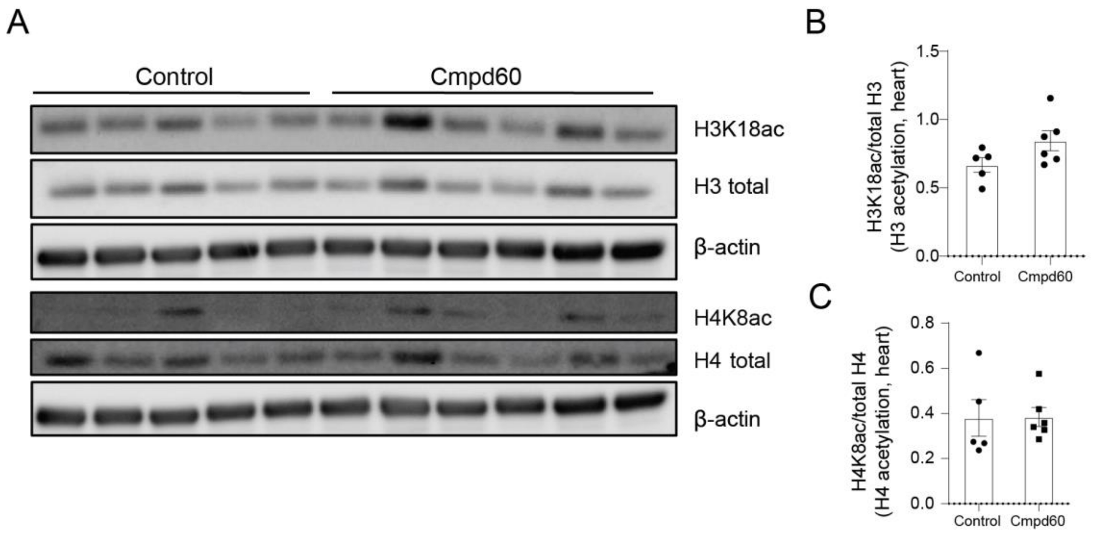


**Supplemental Figure 4 | The effect of Cmpd60 on heart tissue**

A) Representative western blot showing expression of H3K18ac, total H3, H4K8ac, H4 total and β-Actin in aged hearts treated with control or Cmpd60 as indicated in Figure 2B (n=5-6/group). B) Relative expression of Histone H3 acetylation levels (H3k8Ac) assessed by western blot in heart tissue of aged mice treated with control and Cmpd60. Protein expression was normalized against H3 total and expressed as mean ±SEM (n=6/group). C) Relative expression of Histone H4 acetylation levels (H4k8Ac) assessed by western blot in brain tissue of aged mice treated with control and Cmpd60. Protein expression was normalized against H4 total and expressed as mean ±SEM (n=5-6/group).
